# Muscle torques provide more sensitive measures of post-stroke movement deficits than joint angles

**DOI:** 10.1101/642272

**Authors:** Ariel B. Thomas, Erienne V. Olesh, Amelia Adcock, Valeriya Gritsenko

## Abstract

The whole repertoire of complex human motion is enabled by forces applied by our muscles and controlled by the nervous system. The of stroke on the complex multi-joint motor control is difficult to quantify in a meaningful way that informs about the underlying deficit in the active motor control and intersegmental coordination. We tested the idea that post-stroke deficit can be quantified with high sensitivity using motion capture and inverse modeling of a broad range of reaching movements. Our hypothesis is that muscle moments estimated based on active joint torques provide a more sensitive measure of post-stroke motor deficits than joint angle and angular velocity. The motion of twenty-two participants was captured while performing reaching movements in a center-out task, presented in virtual reality. We used inverse dynamics analysis to derive active joint torques that were the result of muscle contractions, termed muscle torques, that caused the recorded multi-joint motion. We then applied a novel analysis to separate the component of muscle torque related to gravity compensation from that related to intersegmental dynamics. Our results show that individual reaching movements can be characterized with higher information content using muscle torques rather than joint angles. Moreover, muscle torques allow for distinguishing between the individual motor deficits due to aging or stroke from the typical differences in reaching between healthy individuals. This novel quantitative assessment method may be used in conjunction with home-based gaming motion-capture technology for remote monitoring of motor deficits and inform the development of evidence-based robotic therapy interventions.

**New and Noteworthy:** Functional deficits seen in task performance have biomechanical underpinnings, seen only through the analysis of forces. Our study has shown that estimating muscle moments can quantify with high sensitivity post-stroke deficits in intersegmental coordination. An assessment developed based on this method could help quantify less observable deficits in mildly affected stroke patients. It may also bridge the gap between evidence from studies of constrained or robotically manipulated movements and research with functional and unconstrained movements.

## Introduction

Movement is a complex interplay between forces generated by our muscles under the control of the central nervous system and the environment. Neurological diseases such as stroke damage the neuromuscular mechanisms of movement production. The resulting movement deficits are a major contributing factor to stroke being the leading cause of long-term disability in the United States (Benjamin et al. 2019). The clinical assessment of movement deficits is based on the observation of movements by experts. For example, sophisticated clinical tests such as the Fugl-Meyer Test of impairment (Fugl-Meyer et al. 1974) or the Wolf Motor Function Test and the Action Research Arm tests of functional ability (van der Lee et al. 2001; Wolf et al. 2001) rely on experts timing and rating the quality of several observed movements, some of which are focused on moving single joints while others on complex functional tasks. However, these tests cannot account for the redundancy of the arm anatomy that underlie inter-subject variability, i.e. the typical differences in how movements are produced by individuals. For example, the same endpoint movement can be performed with different combinations of peak shoulder flexion and abduction angles, i.e., with more or less elbow elevation, which in turn affects the elbow angle. The challenge for motor assessment is in distinguishing the typical variability due to this redundancy from the abnormal changes in movement caused by stroke. Furthermore, the prevalence of stroke increases with age (Virani Salim S. et al. 2021) so that sarcopenia and age-related neurological changes further contribute to the individual differences in how movements are produced (Dutta and Hadley 1995; Edström et al. 2007; Evans 1995; Evans and Campbell 1993; Seidler et al. 2010; Shaffer and Harrison 2007). To compensate for this individual variability in how movements are generated, clinical tests typically employ low-resolution scoring system and extensive training of raters to maximize inter-rater validity. However, this leads to reduced responsiveness and predictive validity of clinical tests and ceiling effects in patients with mild motor deficits (Hsieh et al. 2009; van der Lee et al. 2001). Importantly, randomized clinical trials show variable effectiveness of treatments developed based on the evidence provided by these clinical assessments (Duncan et al. 2003; Saposnik et al. 2016; Wolf et al. 2010). It is widely believed that higher quality assessment measures for informing the design of future interventions are needed (Krakauer and Carmichael 2017; Pollock et al. 2014).

Motion capture offers an objective way to assess movement deficits using algorithms (Schwarz et al. 2019). Measures derived from motion capture, such as the temporal profiles of joint angle changes during movements, are directly relatable to existing clinical assessments (Chang et al. 2011; Clark et al. 2013; Mousavi Hondori and Khademi 2014; Olesh et al. 2014). Metrics that capture inter-subject and inter-trial variability are preferred as they help to standardize motor assessment based on motion capture (Schwarz et al. 2019). In our study, we were inspired by these recommendations and used the performance index, a measure that was originally developed by Fitts to quantify speed-accuracy tradeoff (Fitts 1954). The performance index quantifies the information content of movement trajectories based on amplitude, accuracy, and timing of movement, combining the different variables that contribute to inter-subject and intertrial variability. We have supplemented this analysis with a direct comparison of the temporal profiles of kinematic and dynamic variables between limbs. This metric derived from joint angles is closely related to the qualitative scores of Fugl-Meyer Assessment (Olesh et al. 2014). Here we used both the performance index and the coefficient of determination derived from joint angle and muscle torque trajectories to compare their relative information content and gain insight into the potential of these metrics to assess age-related and post-stroke motor deficits.

The 2019 Stroke Recovery and Rehabilitation Roundtable recommended post-stroke movement analysis to include motion capture of planar (2D) reaching tasks with gravitational support and unsupported three-dimensional (3D) functional tasks (Kwakkel et al. 2019). The functional tasks were recommended specifically to enable quantifying intersegmental or intersegmental coordination using the analysis of natural, unrestrained, and simultaneous movement of multiple joints (Schwarz et al. 2019). Moreover, the analysis of joint angles as independent degrees of freedom (DOFs) enables the identification of compensatory strategies, when for example increased shoulder abduction helps to compensate for reduced elbow mobility (Murphy et al. 2011). In our study we applied similar analyses to study intersegmental coordination in several people with chronic stroke and compare it to the intersegmental coordination in young and aged controls. However, measuring kinematic variables, such as joint angles, can provide only indirect information about how the movement is generated by neuromuscular action. It is not possible to infer from kinematic measures what forces were applied by the muscles to make the arm move, even for a seemingly simple but clinically important determination of whether the motion is active or passive. The forces muscles produce can be estimated by inverse modeling using the equations of motion. The temporal profiles of joint angles obtained from motion capture are differentiated, combined with the estimates of segment inertia, and plugged into the equations of motion to compute net, passive, and active or applied joint torques (Ellis et al. 2009; Russo et al. 2014; Sainburg et al. 1995; Shabbott and Sainburg 2008). This inverse modeling approach takes into account passive torques caused by complex limb inertia and extrinsic forces, such as gravity or objects held in the hand. The active joint torques, termed muscle torques, are the summed result of all the moments generated by muscle contractions representing only the applied forces necessary to produce the observed motion in presence of all the internal and external passive forces (Dounskaia and Wang 2014; Gentili et al. 2007; Le Seac’h and McIntyre 2007; Papaxanthis et al. 2005; Wang and Dounskaia 2016). Muscle torques can then be used to quantify the post-stroke changes in the production of active movement while controlling for the role of passive forces.

The component of muscle torque that is responsible for intersegmental coordination can be extracted computationally. The inverse dynamic simulations used to estimate muscle torque can be ran without gravity, simulating the active torques necessary to produce the observed motion without gravity (Olesh et al. 2017). This dynamic component of muscle torque also reflects the forces that are produced during planar reaches with gravitational support. Such 2D movements are used for quantifying post-stroke motor deficits and robot-assisted rehabilitation (Coderre et al. 2010; Keeling et al. 2021; Kwakkel et al. 2019; Scott and Norman 2003). Studies of multisegmented arm motion with and without gravity have shown that this dynamic component of muscle torque reflects intersegmental coordination (Debicki and Gribble 2005; Gribble and Ostry 1999; Ketcham et al. 2004; Pigeon et al. 2003). Deficits in intersegmental coordination have been reported in people with cerebellar strokes (Bastian et al. 1996). The dynamic component of muscle torque is thought to be related to the phasic component of muscle activity, which is the remainder after subtraction of the tonic posture-related component of muscle activity (Olesh et al. 2017). Moreover, it has been shown that cerebellar damage preferentially impairs phasic EMG (Manto and Bosse 2003). Therefore, quantifying the dynamic component of muscle torque may provide a method to assess post-stroke disruption in intersegmental coordination.

Our study aim was to quantify objectively post-stroke motor deficits using motion capture and inverse dynamics of stereotypical reaching movements. Here we define a motor deficit narrowly as a statistical difference in individual metric derived from kinematic or dynamic variable of movement from the mean metric for young controls. Our hypothesis is that muscle torques contain more information about the post-stroke motor deficits than angular kinematics. Because of the small sample size and the heterogeneity of our sample, we designed our experiment to quantify individual motor deficits in participants with stroke relative to aged individuals and the group mean of young controls. We then applied a novel analysis that separated muscle torques into components related to gravity compensation and intersegmental coordination and used these and kinematic measures to quantify motor deficits.

## Methods

### Participants

Twenty-two human participants were recruited to perform reaching movements to virtual targets with both arms. The participants were divided into three groups, Control, Stroke, and Aged. The Control group (23 ± 1.2 years) included 9 participants without any known neurological or musculoskeletal disorders. The kinematic and torque data from the Control group were reported in Olesh et al. (2017). The Stroke group included 8 participants (58 ± 6.9 years), who have suffered single unilateral ischemic stroke at least three months prior to the experiment (Table 1). Diagnosis was performed by neurologists during hospital admission or following neurological evaluations at Ruby Memorial Hospital. Individuals were excluded if they could not produce visible movement with their shoulder and elbow, or if they were unable to provide written consent to participate. The Aged group (58 ± 2.4 years) included 5 participants without any known neurological or musculoskeletal disorders, whose age was within the mean ± SD of participants in the Stroke group. All control participants were right-hand dominant and reported no unrelated movement disorders or significant injuries to their upper extremities. The study and the consent procedure were approved by the Institutional Review Board of West Virginia University (Protocol # 1311129283). All participants provided written consent before participation.

**Table 1.** Characteristics of individuals with stroke.

| ID | Gender | Infarct hemisphere | Infarct description and location | Age | Years post stroke | Contra-lesional reach duration (s) | Contra-lesional endpoint accuracy (cm) |
| --- | --- | --- | --- | --- | --- | --- | --- |
| S1 | F | Left | Left MCA | 51 | 3 | $0.7 \pm 0.16^*$ | $4.5 \pm 0.87$ |
| S2 | M | Left | Left caudate lenticular nucleus and external horn of the left ventricle | 58 | 5 | $0.8 \pm 0.11^*$ | $4.1 \pm 1.22$ |
| S3 | M | Right | Right dorsal pontine-medullary lacunar infarction | 67 | 0.5 | $0.7 \pm 0.17^*$ | $5.6 \pm 1.37$ |
| S4 | M | Right | Lacunar infarct in posterior right putamen and border of right internal capsule | 68 | 0.25 | $0.5 \pm 0.08$ | $4.5 \pm 1.04$ |
| S5 | M | Left | Left MCA | 53 | 0.5 | $1.3 \pm 0.52^*$ | $8.2 \pm 2.80$ |
| S6 | M | Right | Right MCA | 60 | 6 | 0.5 ± 0.10 | 4.3 ± 1.37 |
| S7 | M | Right | Right lateral medullary infarction with occluded right vertebral artery | 51 | 8 | 1.4 ± 0.49* | 8.6 ± 2.63 |
| S8 | M | Right | Right MCA, extending posteriorly | 53 | 7 | 0.4 ± 0.12 | 7.2 ± 3.30 |

MCA stands for middle cerebral artery. Standard deviations are across movement directions. \* indicates significant difference from Control group with $p < 0.0031$ for all; Significant alpha with Bonferroni correction for repeated measures = 0.0031. Note, that the amplitude of movement in S8 participant was decreased due to his inability to reach the same distance as controls.

### Experimental Task

During the experiment, participants reached to virtual targets in a center-out task, as described in detail in Olesh et al. (2017) (Fig. 1). The arm was not supported, all movements were unconstrained. Movements were instructed using a virtual reality (VR) software (Vizard by Worldviz) and headset (Oculus Rift), which randomly displayed one of 14 targets, 10 cm in diameter, arranged equidistantly from a center target. A center target was placed in the VR space so that the initial posture of the upper arm was aligned with the trunk (all shoulder angles at 0°) and the forearm parallel to the floor palm down (elbow angle at 90° and wrist at 0°; Fig. 1A). This minimized both the intertrial variability and inter-subject variability in motion trajectories. For each participant, the distance from the center target to the peripheral targets was scaled to 30% of arm length (anterior acromial point to the distal tip of the index finger). This distance between the central and peripheral targets was on average 20 cm. One most impaired participant (S8, Table 1) was unable to reach targets reliably with his contralesional arm. Therefore, the reaching distance was decreased to 10 cm only for the contralesional arm.

**Figure 1.**
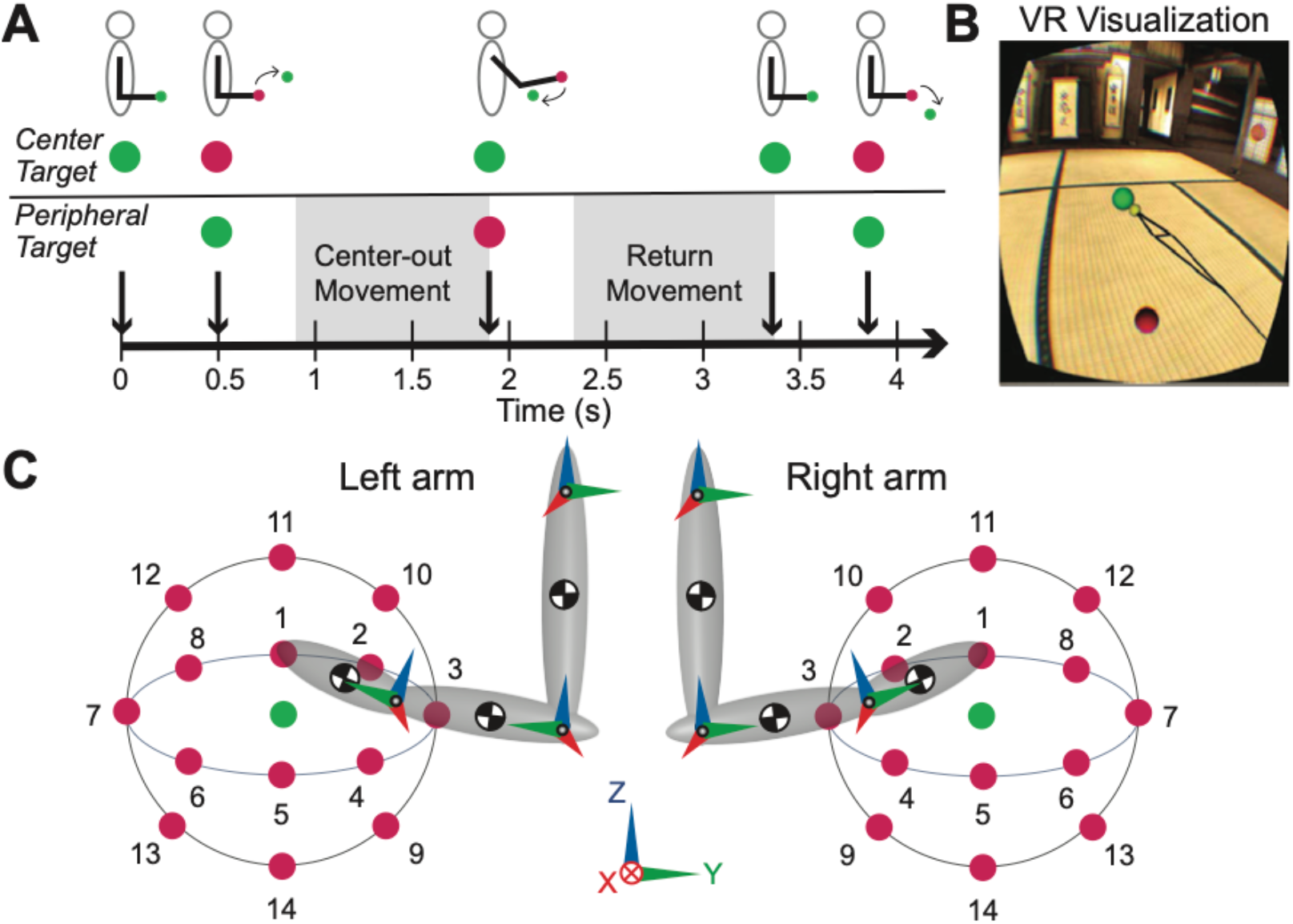
Standardized reaching in virtual reality. **A.** Trial timeline for each center-out and return reaching movement toward one of 14 virtual targets. **B.** Participant’s VR view of the targets with visual feedback of their limb position (black “stick figure” indicating hand, forearm, and upper arm), the yellow sphere indicated the position of the tip of the index finger. **C.** Side view of the scaled dynamic models of the right and left arms and relative target locations. The checkered circles represent the centers of mass of the segments. Arrows represent the orientations of local and world coordinate systems used for defining joint angles and the degrees of freedom for the right and left arms. Shoulder, elbow, and wrist flexion/extension degrees of freedom were calculated around X (red) axes. Shoulder abduction/adduction degree of freedom was calculated around Y (green) axis. Shoulder internal/external rotation degree of freedom was calculated around Z (blue) axis.

Participants were seated and instructed to reach to targets as quickly and as accurately as possible without moving their torso. The torso motion was restrained with Velcro straps attached to the vertical backrest of a chair. The participant’s arm was visualized in VR as a “stick figure” connecting the positions of motion capture markers (Fig. 1B) and it moved concurrently with the participant at the temporal resolution of the VR helmet refresh rate (~90 Hz). Index fingertip was shown as a yellow ball 5 cm in diameter, participants were instructed to move it into the center of each target. Individual joint motion of the digits was not tracked; therefore, all participants were instructed to keep palm flat with all fingers extended and wrist pronated (palm down). The arm was not supported during reaching. Participants with stroke wore a finger splint made of polystyrene foam weighing < 10 gram to keep the digits 2-5 extended including at the metacarpophalangeal joints (palm down and open).

Each movement began with the participant's hand in the center target. A movement was cued by the center target changing color from green to red and the appearance of one peripheral green target (Fig. 1B). Upon the detection of index fingertip inside the peripheral target radius, its color changed to red, cueing the participant to return to the central target (Fig. 1A, end of center-out movement). After the participant reached the center target (Fig. 1A, end of return movement), the task reset, the peripheral target disappeared, and a new one appeared after a half-second delay. Movements to each peripheral target location were performed in a randomized order and repeated 15 times with rest breaks after bouts of 70 trials or upon request. Each participant repeated this experiment with both arms in separate sessions on different days.

### Data Collection and Processing

All analyses and statistics were performed in MATLAB (Mathworks Inc.). Motion capture was recorded during the experiment using an active-marker motion capture system (Impulse by PhaseSpace). The light emitting diodes (active markers, LEDs) were placed according to best practice guidelines on anatomical bony landmarks of the arm and trunk (Robertson et al. 2013). Motion capture was collected at a rate of 480 frames per second, low pass filtered at 10 Hz and interpolated with a cubic spline (maximum interpolated gap: 0.2 s). Data synchronization methods are described in detail in Talkington et al. (2015). To improve the quality of kinematic data, spike artifacts detected at greater than 3 standard deviations at the elbow LED were removed automatically by blanking within .05 seconds of the spike before interpolation. Joint angles were calculated from motion capture using local coordinate systems defined as follows. Six LEDs on the clavicle, sternum, spine and the shoulder of the analyzed arm were used to define the trunk coordinate system. All joint angles were calculated relative to the trunk coordinate system, thus controlling for any change in trunk posture. The orientation of the trunk coordinate system was the same for both arms, therefore the directions of motion about the shoulder flexion/extension and internal/external rotation DOFs were opposite between limbs. They were flipped in the correlative analysis described below. Three LEDs, 2 on the shoulder and 1 on the elbow, were used to define upper arm coordinate system. Three LEDs, 1 on the elbow and 2 on the wrist, were used to define forearm coordinate system. Three LEDs, 2 on the wrist and 1 on the fingertip, were used to define hand coordinate system. The axes of the local coordinate systems were oriented in the same direction for both arms as shown in Fig. 1C. Joint angles were defined as Euler angles that corresponded to five joint degrees of freedom (DOFs) including 3 shoulder DOFs flexion/extension, abduction/adduction, internal/external rotation, 1 elbow DOF flexion/extension, and 1 wrist DOF flexion/extension. In some participants, the medial wrist LED was not reliably tracked due to being obscured frequently from camera view by moving body segments. Therefore, wrist pronation/supination DOF was not reliably detected and, thus, excluded from analysis. Wrist abduction/adduction was found to be minimal during these tasks and was likewise not included in the analysis.

The onset and offset of each center-out and return movements were identified from the differentiated trajectory of hand marker (velocity) crossing the threshold of 5% of maximal velocity at the beginning and the end of a given movement. Center-out movements toward one of the peripheral targets were separated from the return movements toward the central target by these events and analyzed as separate movements. The events were verified through visual inspection of the plotted trajectories to correct for unintended or corrective motion around the virtual target. The onset and offset events were used for temporal normalization of kinematic and torque profiles prior to averaging. Profiles starting 100 ms prior to the onset events were included in the analysis of each movement to capture the full profile of phase-advanced torques. All values included in text are means ± standard deviation across participants, unless otherwise indicated.

### Inverse Dynamics

The arm model was implemented in the Multibody toolbox of MATLAB. The model with 5 DOFs described in Olesh et al. (2017) was used to calculate forces at the shoulder, elbow, and wrist joints. Joint angles for the shoulder, elbow, and wrist, obtained as described above were used to drive the model and simulate the center-out and return movements. The trunk was assumed to be stationary and in line with the world coordinate system (Fig. 1C). The inertia of major limb segments (humerus, radius/ulna, and hand) were modeled as ellipsoids with the long axes and masses scaled to the lengths of individual participants (Winter 2009). The segment diameters were kept constant at 10 cm, 6 cm, and 6 cm for upper-arm, forearm, and hand respectively. The short axes of the ellipsoids remained constant across participants. Inverse dynamics simulations using the individually scaled model shown in Fig. 1C were ran in Simulink. The output of these simulations was applied torques, not to be confused with net torques commonly referred to in Newton laws, i.e., τ = I*α, where τ is net toque, I is inertia, and α is angular acceleration. The applied torques (τ_*a*_) represent the active torques that need to be applied externally, in our case by muscles, to produce the desired motion of the limb in presence of passive torques (τ_*p*_) generated by the inertia and external forces, such as gravity. Torques summate, thus τ = τ_*a*_+ τ_*p*_ or τ_*a*_ = τ − τ_*p*_. We termed the applied torques muscle torques (MT). MT were further subdivided into two additive components, termed gravitational and dynamic components. The gravitational component (MTg) captured the portion of applied torque that supports the limb segments against the force of gravity. The dynamic component (MTd) captured the residual applied torque related to intersegmental coordination (Gottlieb et al. 1997; Russo et al. 2014). To obtain MTd, we ran the inverse dynamics simulations with gravity of the physics engine set to zero (Olesh et al. 2017). As before, we took advantage of the additive nature of torques, deriving MTg by subtracting applied torques calculated in simulations without gravity from those calculated in simulations with gravity, i.e. MTg = MT – MTd, for each DOF movement and subject.

The quality of these simulations was checked by running forward simulations and calculating the root-mean-squared error (RMSE) between the endpoint trajectories obtained from motion capture and those from forward simulations. The RMSE mean ± standard deviation across participants for Control group was 0.0014 ± 0.0020 and 0.00014 ± 0.00035 mm for the right and left arm respectively, for the Aged group it was 0.00027 ± 0.00046 and 4.5e-05 ± 4.7e-05 mm for the right and left arm respectively, and for the Stroke group it was 0.0049 ± 0.0072mm and 0.3 ± 0.8 mm for the ipsilesional and contralesional arm respectively. The RMSE values in the Stroke group were larger due to lower quality of motion capture data, where the markers disappeared from the view of cameras more often than during experiments with participants in other groups.

The movement time and endpoint accuracy were calculated for each participant. Endpoint accuracy was calculated as fingertip distance to the center of the outer targets at the end of the center-out reach phase, described above. Movement time was calculated as the elapsed time between the onset and offset events of each forward and return movement.

### Metrics

Angular kinematic and torque profiles were normalized in time using onsets and offsets of each movement and resampled (1000 samples). Intertrial variability was calculated as the standard deviation of repetitions of individual movements toward the same target (n = 15) across the normalized trajectory, and then averaged over time. The inter-trial variability for wrist joint angle in young controls was 5.3 ± 0.11 and 5.7 ± 0.16 degrees for the right and left arm respectively. This was larger than what was observed in the moving DOFs of the shoulder (F/E R: 3.2 ± 0.3, L: 3.4 ± 0.36; Ad/Ab R: 3.4 ± 0.43, L: 2.9 ± 0.33; InR/ExR R: 3.7 ± 0.45, L: 3.9 ± 0.45 degrees) and elbow (L: 3.8 ± 0.5, R: 4.3 ± 0.11 degrees). Therefore, wrist DOF was removed from the rest of analysis. The individual normalized trajectories for shoulder and elbow were averaged to create a mean profile for each center-out and return movement toward each target for each participant. Muscle torque and muscle torque components were normalized by the subject-specific weight of each arm segment used during the inverse dynamic simulations, allowing for comparisons between subjects. From these, two types of metrics were derived, performance index and coefficient of determination

Performance index (*I*) was calculated from the time-normalized trajectories of joint angles, angular velocities, muscle torques (MT) and their components (MTd and MTg) using the following formula adapted from (Fitts 1954) for each center-out or return movement to or from a given target per DOF of right or left arm for each subject:

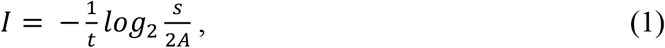

where *t* is the mean time to perform a given movement; *A* is the peak-to-peak amplitude of a mean time-normalized trajectory for a given DOF, movement, and subject, *s* is the standard deviation of *A* across repetitions of center-out or return movement. The units of the performance index are bits/s.

The coefficient of determination (R^2^) was calculated between the time-normalized trajectories of joint angles, angular velocities, muscle torques (MT) and their components (MTd and MTg) for corresponding signals from the right and left limbs of the same participant. In this analysis, the participants served as their own controls, which reduced inter-subject variability. Mirror movements produced by left and right arms were matched as numbered in Fig. 1C because they were produced with the same joint angles and required contractions of the same muscles to perform them. R^2^ ranges between 0, indicating low symmetry between trajectories of right and left limb, and 1 indicating highly symmetrical trajectories.

### Statistical Analysis

Data in Results are reported as means ± standard deviation either across participants within each group or across movement directions for each individual, unless otherwise described. Paired t-tests were applied to all metrics comparing mean values across controls per movement direction and the corresponding data from the matching side in individuals in the Aged and/or Stroke groups. Familywise error was corrected for multiple testing using Bonferroni method, i.e. α /N, where N is the number of subjects in either Aged or Stroke group. The significant α is reported in Results with every statistic.

To compare the sensitivity of distinguishing post-stroke motor deficits from inter-subject variability based on joint angles and MTd, we used k-means clustering to classify participants into one of 2 groups, those with motor deficits and those without motor deficits. Clustering was applied to either angular or MTd R^2^ for each DOF of the shoulder and elbow per center-out and return movements per subject. To identify clusters, we used squared Euclidean distance measure with a heuristic approach for cluster center initialization (Arthur and Vassilvitskii 2007). The reproducibility of cluster assignment was tested by running the algorithm 100 times and reporting the chances of aged individuals being classified into the same cluster as young controls and the chances of individuals with stroke being classified into a separate cluster from that with young controls.

Supplementary materials and original data are included in the GitHub repository https://github.com/NeuroRehabLab/Stroke_Dynamic_Scoring

## Results

All participants in the Control group performed the center-out and return reaching movements with low inter-subject and inter-limb variability in endpoint trajectories (Fig. 2A). The reaches were performed with preferred speed, which differed somewhat between individuals. The endpoint accuracy defined as the distance from the center of the virtual target to the end of endpoint trajectory was 3.69 ± 0.9 cm for right and 3.72 ± 0.5 cm for left arm, well within the required tolerance. Joint angle trajectories mirrored the sigmoidal shape of the endpoint trajectories with classical bell-shaped velocity profiles but contained more detailed information about the motion of individual joints. Because movement amplitude, accuracy, and variability are inter-related, we quantified the performance of individuals using information theory as described in methods and earlier publications (Fitts 1954; Young et al. 2009). The performance index based on joint angles was 4.7 ± 1.5 bits/s (mean ± standard deviation across individuals) for the right arm and 4.4 ± 1.2 bits/s for the left arm reaches. These values are much lower than values close to 10 bits/s reported for ballistic movements (Fitts 1954), indicating that the reaching movements in our study were not challenging. Similar amount of information was reflected in the MTg trajectories (3.6 ± 1.2 and 3.6 ± 1.0 bits/s for right and left arm respectively), likely because this component is related to the cosine of the orientation of the segments to gravity, i.e., it is a dependent variable of joint angle. In contrast, higher information content was reflected in the trajectories of other variables. The performance index based on angular velocity was 5.8 ± 1.6 and 5.3 ± 1.2 bits/s for right and left arms respectively, and the performance index based MTd was 6.3 ± 2.0 and 5.7 ± 1.4 bits/s for right and left arms respectively. As defined in Methods, muscle torque is the sum of MTg and MTd. Therefore, the MT performance index was intermediate between that derived from the two components at 4.0 ± 1.6 and 4.1 ± 1.3 bits/s for right and left arms respectively. The difference between the MTd and angular performance indices was 1.7 ± 0.9 for the right arm and 1.3 ± 0.3 for the left arm. This shows that the performance of reaching movements by the participants in the Control group can be characterized with higher information content using MTd trajectories compared to that using trajectories of joint angles.

**Figure 2.**
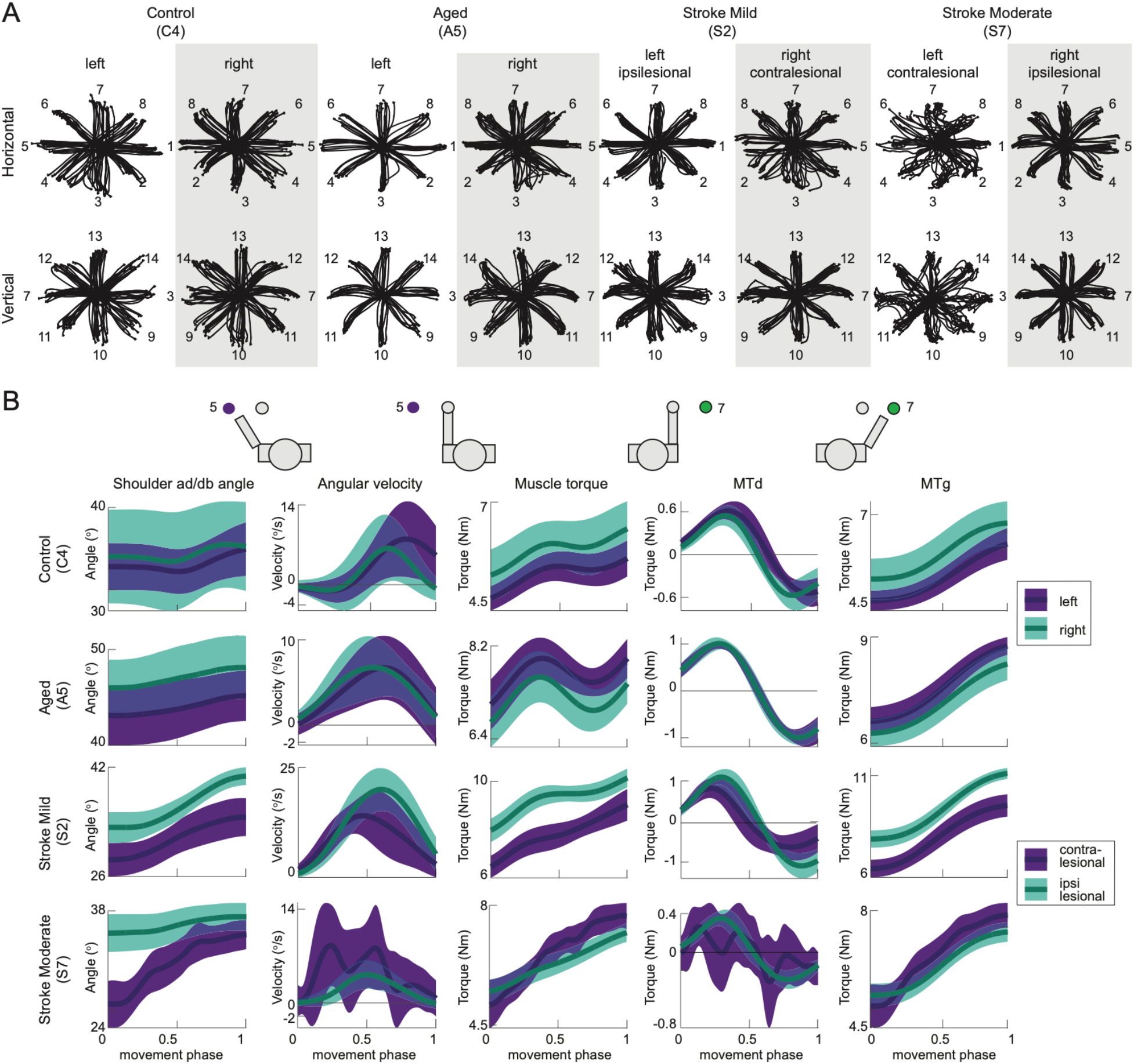
Kinematic and torque profiles of several participants. **A.** Endpoint trajectories for individual center-out reaching movement toward each of 14 peripheral targets. **B.** Example mean kinematic and dynamic trajectories from one movement, toward target 5 for left arm and toward target 7 for right arm. Mean trajectories (lines) and standard deviations (shaded areas) are shown for shoulder abduction/adduction DOF.

As expected, all five participants in the Aged group were able to perform the center-out and return movements with matched reach amplitudes and targets positions (Fig. 2A). The reaches were slower in 3 out of 5 aged participants compared to that for the corresponding arm of the Control group participants (A1 right arm (R): t = 4.1, p = 0.0004; A3 R: t = 8.7, p < 0.0001; A4 R: t = 13.1, p < 0.0001; and A1 left arm reach (L): t = 3.8, p = 0.0007, significant α = 0.0042). The endpoint accuracy of most aged participants was the same as in Controls, with the exception of A2, whose accuracy was worse than in controls (R: t = −6.3, p < 0.0001 and L: t = −6.3, p < 0.0038 for all). This shows that most participants in the Aged group reached slower to maintain the accuracy specified by the virtual targets, demonstrating the classical speed-accuracy tradeoff (Fitts 1954; Young et al. 2009). The angular performance index in aged participants was on average similar to young controls at 4.1 ± 1.3 bits/s for the right arm and 4.0 ± 0.9 bits/s for the left arm reaches. As in the Control group, the MTd performance index was the largest among all indices at 5.1 ± 1.6 bits/s for the right arm and 5.9 ± 0.8 bits/s for the left arm reaches. The difference between the MTd and angular performance index was 1.0 ± 0.4 bits/s for the right arm and 1.9 ± 0.4 bits/s for the left arm. The angular performance index of all aged participants was different from that of Control group for reaching with one arm and sometimes both (A1 R: t = 10, L: t = 6; A2 L: t = 6; A3 R: t = 6, L: t = −4; A4 R: t = 7 and A5 R: t = −5) with all p-values less than α = 0.0042 with Bonferroni correction for familywise error. The MTd performance index of aged participants was also higher that the angular performance index, just as was observed for the Control group. This increase was meaningful, as it showed larger changes in information content of the same aged participants (A1 R: t = 16, L: t = 6; A2 R: t = 7, L: t = 8; A3 R: t = 15, L: t = −7; A4 R: t = 15 and A5 R: t = −5, L: t = −6, p < 0.0042) as those identified by the angular performance index. This shows that the algorithm based on MTd trajectories is the most sensitive to individual differences in reaching. Importantly, this shows that all aged participants reached differently from young controls with at least one arm, suggesting a propensity for asymmetric movement in the elderly.

All eight participants with stroke were able to perform the center-out and return movements with matched reach amplitudes and targets positions using their ipsilesional arm. The reaches were slower in 6 out of 8 stroke participants compared to that for the corresponding arm of controls (S1: t = 3.9, S3: t = 13.3, S5: t = 16.1, S6: t = 5.9, S7: t = 21.1 and S8: t = 11.8, p was less than the significant α = 0.0031 for all). The endpoint accuracy of reaching with the ipsilesional arm was similar to the accuracy of reaching with the corresponding arm in Controls likely due to the speed-accuracy tradeoff (Fitts 1954; Young et al. 2009). The angular performance index for the ipsilesional arm was on average lower than that for the corresponding arm of aged and young controls at 3.3 ± 1.2 bits/s. As in the controls, the MTd performance index was the largest among all indices at 4.1 ± 1.4 bits/s for the ipsilesional arm with a difference between them at 0.9 ± 0.4 bits/s. The angular performance index for the ipsilesional arm was significantly different form aged controls in 4 participants with stroke (S4 t = −5, S5 t = 4, S7 t = 11, and S8 t = 8, p < 0.0031 for all). These differences in S7 and S8 were larger when compared to the corresponding mean performance index of the Control group (data not shown). The differences in MTd performance index for the ipsilateral arm were significant for 5 participants with stroke (S3 t = 5, S4 t = −5, S5 t = 8, S7 t = 19 and S8 t = 13, p < 0.0031 for all), only some of them moved slower. The same was true for the comparison with the Control group (data not shown). Once again, the sensitivity of the algorithm based on MTd trajectories was most sensitive to the individual differences in reaching compared to that based on trajectories of angles. Overall, this shows that the changes in reaching with the ipsilesional arm in 5 (S3, S4, S5, S7 & S8) out of 8 participants with stroke were different from the changes in reaching associated with age and different from reaching by young controls. In the rest of participants with stroke (S1, S2 and S6) reaching with the ipsilesional arm was similar to that of aged or young controls.

All but two participants in the Stroke group were able to perform the center-out and return movements with matched reach amplitudes and targets positions using their contralesional arm (Fig. 2A). The movement speed of reaching with the contralesional ‘impaired’ arm was lower than that of Control group (Table 1). The contralesional reach durations were longer in all but three participants with stroke compared to that for the matching arm in Control group (Table 1; S1 t = 6.7, S2 t = 9.0, S3 t = 9.9, S5 t = 10.3, and S7 t = 17.0, p < 0.0031 for all). However, the endpoint accuracy of reaching with contralesional arm was similar to that of the Control group in all participants with stroke due to the speed-accuracy tradeoff (Table 1). Of note is that the timing and accuracy measures are confounded by the shorter reach distances of one participant (S8). The performance index controls for these confounding factors. The angular performance index for the contralesional arm was on average lower than that for the corresponding arm of aged and young controls at 3.2 ± 1.2 bits/s. As in the controls, the MTd performance index was the largest among all indices at 4.7 ± 1.7 bits/s for the contralesional arm with a difference between them at 1.5 ± 1.4 bits/s. Angular performance index for the contralesional arm was lower in all but 2 participants with stroke compared to that for the corresponding arm of participants in the Aged group (S2 t = 6, S3 t = 6, S4 t = −4, S5 t = 10, S7 t = 12 and S8 t = 4, p < 0.0031 for all). Comparison to the Control group revealed similar differences (data not shown). Moreover, MTd performance index revealed larger positive differences in S1 and in most participants with significant differences in their angular performance index (S1 t = 6, S2 t = 24, S3 t = 18, S4 t = −3, S5 t = 34 and S7 t = 41, p < 0.0031 for all) with the exception of S8. Comparison with the Control group revealed similar differences (Fig. 3). In conclusion, altogether these metrics have shown that reaching with the contralesional arm in most participants was different from that in aged and young controls with the exception of S6.

**Figure 3.**
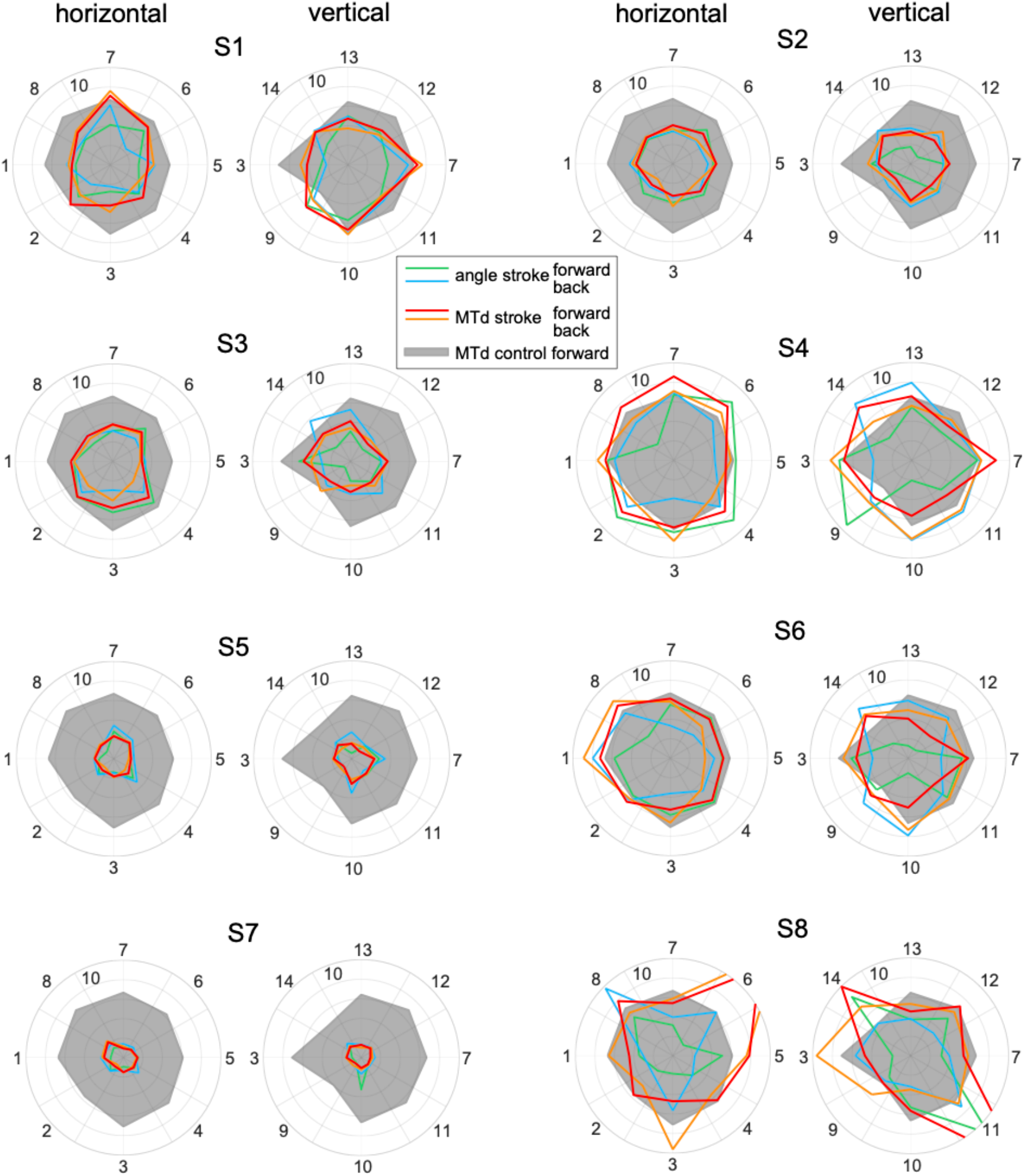
Performance index for each movement direction per participant with stroke. Each plot shows angular performance index (blue and green lines for center-out and return movements respectively) and MTd performance index (red and orange lines for center-out and return movements respectively) for reaching with the contralesional arm. Black shaded area denotes the mean MTd performance index for center-out reaches by young controls. The outer radius in all plots is equal to 10 bits/s.

The MTd performance index was similar across movement directions for both outward and return movements in young controls (Fig. 3, grey shading). In contrast, the angular performance index was not uniformly distributed across movement directions, being close to zero for center-out movements toward targets 4 (toward the body and to the right), 9 (toward the body and down), and 10 (toward the body and up, data not shown). However, in the participants with stroke the performance index for their contralesional arm was uniformly lower than in controls for some (S2, S3, S5 and S7) and non-uniformly distributed across movements for others (S1, S4, S6 and S8; Fig. 3). This illustrates the heterogeneity of individual stroke pathologies. The amplitudes of the angular and MTd performance indices also varied independently across different movement directions, indicating that the changes in the angular excursions for movement in different directions are not always accompanied by pronounced changes in the amplitude of joint torques. The converse is also true, so that in some movements during which some joint angles do not change do required different amplitudes of joint torques. Thus, angular and torque trajectories convey fundamentally different information about movement deficits.

According to Eq. (1), the reduction in the performance index can be driven by the increased time to make the movements, increased trajectory amplitude, and decreased inter-trial variability. The latter can be excluded from reducing the performance index as inter-trial variability was increased rather than decreased in participants with stroke (data not shown). Also, the amplitudes of joint angle trajectories were constrained to be the same by our VR task across all but S8 participant, who did not show uniform reduction of his performance indices (Fig. 3, bottom right corner). This constraint also helps to exclude the amplitude of movement from causing the reduction of the performance index. Therefore, for S3, S5, and S7 the decrease in the performance index for the contralesional arm was driven by slower reaching movements. For the rest of the participants with stroke (S1, S2, S4, and S8), the changes in the performance index were driven by the changes in the ratio between inter-trial variability and the trajectory amplitude. Note, that S6 showed no changes in performance index for either arm.

It is difficult to distinguish the mild motor deficits caused by stroke or aging from the normal variability in how individuals make movements. Therefore, we next quantified how symmetrical the reaching trajectories were between the right and left limb using the coefficient of determination (R^2^) that controls for inter-subject variability. The profiles of joint angles were highly symmetrical between limbs in young and some aged controls and less symmetrical in most participants with stroke (Fig. 4A). The angle R^2^ values tended to saturate at 1 likely due to the lower information content of angular trajectories compared to the MTd trajectories. In contrast, the profiles of MTd were less symmetrical in young controls so that the coefficient of determination was more variable between individuals and less likely to saturate (Fig. 4A). Moreover, the differences in MTd R^2^ values between aged and young participants were larger in more movements than the corresponding differences in angle R^2^ (Fig. 4B). Specifically, for aged participants A3 and A4 the profiles of MTd were significantly less symmetrical than in young controls (paired t-tests between mean R^2^ per movement direction in young controls and A3 t = 5 and A4 t = 3, p < significant α of 0.0031 for both). This supports the propensity for asymmetric movement in these aged participants identified with the unilateral changes in performance index. However, the bilateral changes in performance index in aged participants A1, A2, and A5 were not accompanied by the asymmetric MTd profiles, indicating that in these aged participants the bilateral age-related changes in reaching movements were similar in both limbs. Overall, similar to the conclusions from performance analysis, the differences between young controls and aged individuals were greatly amplified when quantified with the coefficient of determination based on MTd.

**Figure 4.**
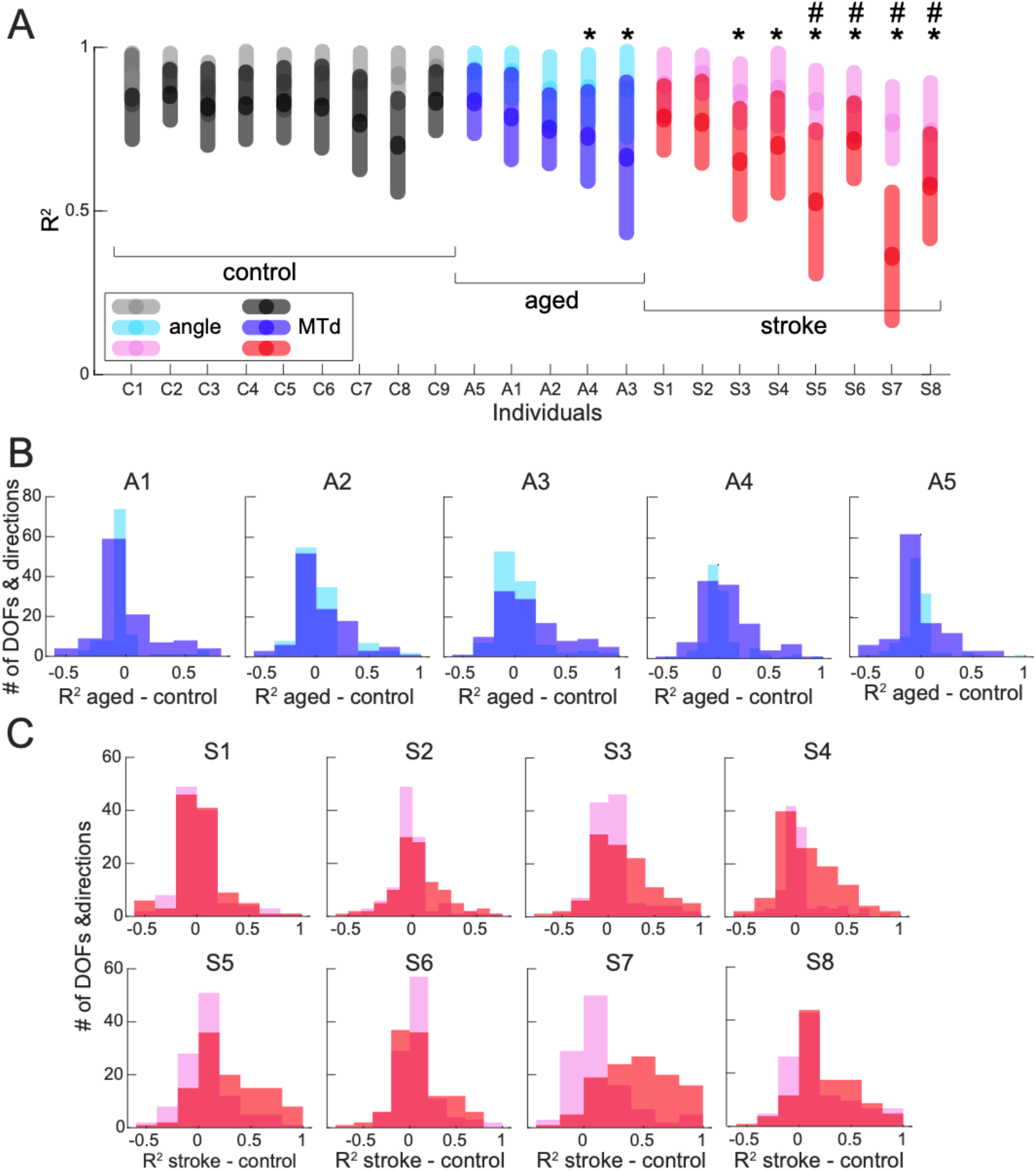
Coefficient of determination (R^2^). **A.** Individual R^2^ based on joint angle and MTd. The central tendencies of data per participant are shown as means (dark circles) with standard deviation across movement directions (light bars). DOFs were averaged per movement direction. Hashtags shows significant differences between angle R^2^ in individuals with stroke compared to angle R^2^ averaged across young controls per movement direction and DOF. Stars shows significant differences between MTd R^2^ in aged individuals and those with stroke compared to MTd R^2^ averaged across young controls per movement direction and DOF. Significant α = 0.0031 with correction for multiple tests. B. Differences between R^2^ in aged and young controls per movement direction per DOF. Colors are as in A. C. Differences between R^2^ in individuals with stroke and young controls per movement direction per DOF. Colors are as in A.

For most participants with stroke, the profiles of both angle and MTd were less symmetrical than the mean R^2^ in young controls (Fig. 4B). For four participants with stroke the angle R^2^ was less symmetrical than the corresponding values in young controls (S5 t = 3, S6 t = 4, S7 t = 6 and S8 t = 6, p < significant α of 0.0031 for all). Furthermore, for six participants with stroke the MTd R^2^ was less symmetrical than the corresponding mean R^2^ in young controls (S3 t = 5, S4 t = 4, S5 t = 10, S6 t = 4 , S7 t = 17, and S8 t = 9, p < significant α of 0.0031 for all). This shows again that the differences between young controls and most individuals with stroke were amplified when quantified with the coefficient of determination based on MTd with the exception of S6. Note, that the performance index of S6 was not different from aged nor young controls, therefore the interlimb asymmetricity of his reaching likely indicates mild hemiparesis. Interestingly, the performance index for S1 and S2 indicated significant differences in reaching with the contralesional arm but not ipsilesional arm. However, the R^2^ values for these participants were not significantly different from young controls, indicating that the shapes of angular and MTd trajectory waveforms were on average symmetrical between limbs. The differences in performance index in these participants were driven by higher inter-trial variability, however the high symmetricity of their reaching trajectories indicate that they do not have pronounced deficits in intersegmental coordination. In contrast, the rest of participants had both significant performance index changes for their contralesional arm driven by both slower movements and increased variability and they also had lower interlimb symmetry in their MTd profiles, indicating larger deficits in intersegmental coordination of contralesional arm.

To further distinguish between motor deficits caused by stroke from changes in movements due to inter-subject variability, the MTd and angle R^2^ values were used to classify individual participants into two groups. K-means clustering analysis relies on an optimization algorithm, which can converge on different solutions. Therefore, the clustering was repeated 100 times. The distance between the two clusters into which all participants were grouped was larger when the analysis was based on MTd R^2^ at 2.94 ± 0.64 across multiple optimizations compared to the analysis based on angle R^2^ at 1.61 ± 0.38 (one-tailed t-test t = 16.35, p = 2•10^−37^). The distances between the two clusters based on MTd R^2^ were consistently larger than those based on angle R^2^ across most movement directions (Fig. 5C). The small distance between clusters based on angle R^2^ values is likely because of these values saturating (Fig. 5A). Consequently, young controls were grouped together 53 – 63 % when clustering was done on angle R^2^ in contrast to 90 – 98 % when clustering was done on MTd R^2^. Similar increases in classification performance were evident for aged participants being grouped together with young controls (angle R^2^: 43 – 64 %, MTd R^2^: 66 – 95 %). Differences in performance indices in participants A2, A3 and A4 and in interlimb asymmetricities in A3 and A4 resulted in these aged participants being grouped less often with young controls based on MTd R^2^ (A2: 82%, A3: 66% and A4: 86% compared to A1: 93% and A5: 95%). This was not as apparent from clustering based on angle R^2^ (A2: 55%, A3: 57% and A4: 43% compared to A1: 64% and A5: 56%). The classification based on MTd R^2^ was also robust for individuals with stroke. The participants with stroke whose motor deficits were minor were rarely separated from controls (S1: 4%, S2: 1%, and S6: 17%), while the participants with stroke whose motor deficits were larger were separated from controls more often (S3: 42%, S4: 27%, S5: 86%, S7: 82%, and S8: 66%). This evidence further supports the conclusion that algorithms based on MTd are more sensitive to subtle differences in motor deficits than those based on joint angle, while still able to separate deficits from typical differences in how movements are produced by individuals.

**Figure 5.**
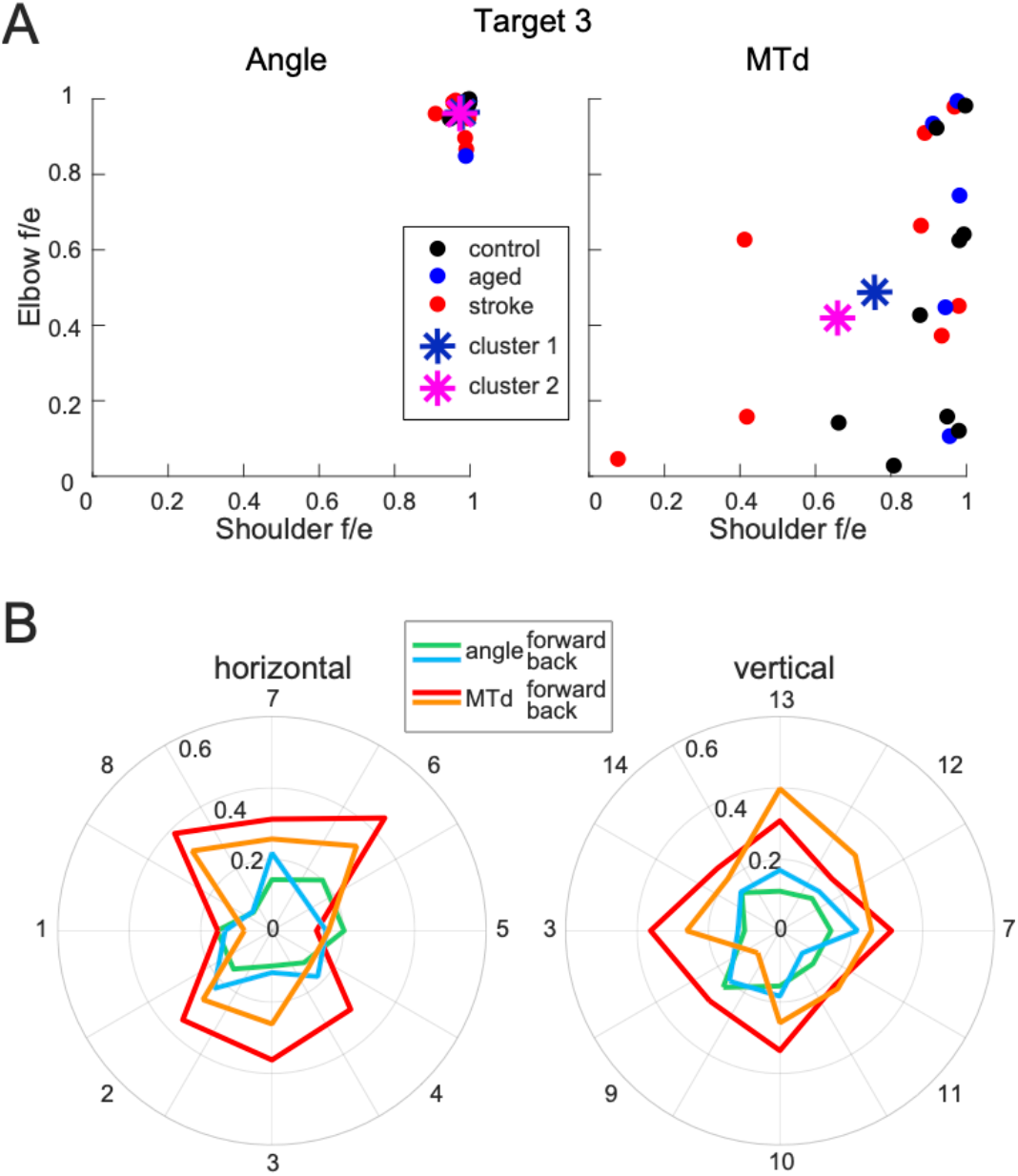
Cluster analysis of the coefficient of determination. **A.** Example cluster coordinates (stars) and individual R^2^ values (filled circles) for a movement toward target 3 for shoulder and elbow flexion/extension (f/e) DOFs. Only 2 DOFs for a single movement are plotted, but the cluster analysis was applied across 112 dimensions of 4 DOFs, 14 center-out, and 14 return movements. Left plot shows results based on angle R^2^ values, while right plot shows results based on MTd R^2^ values **B.** Distances between the two clusters per movement direction. Blue and green lines show distances between clusters of angular R^2^ values for center-out and return movements respectively. Red and orange lines show distances between clusters of MTd R^2^ values for center-out and return movements respectively.

## Discussion

Here we have objectively quantified the individual differences in reaching that could underlie motor deficits caused by aging and stroke. We have shown that joint torques comprise a more sensitive measure of individual differences in reaching compared to joint angles. This confirms our main hypothesis that muscle torques contain more information about the post-stroke motor deficits than angular kinematics. The profiles of the dynamic component of muscle torque in particular contain more information than joint angles do as evidenced by the larger performance index based on the former compared to the latter in all participants. Furthermore, the profiles of the dynamic component of muscle torque were the most sensitive to the individual differences in reaching symmetricity as evidenced by the lowest coefficient of determination values. Furthermore, the clustering analysis has shown that the coefficient of determination based on the dynamic component of muscle torque could be used to distinguish the individual differences in reaching due to aging or stroke from those caused by typical inter-subject variability. A force-based assessment developed based on this method could improve the objective measurement of individual motor impairment. This type of assessment may be especially useful for patients who have less “observable” deficits, such as those classified as asymptomatic via traditional motion-based assessments, but who may still report difficulty moving, increased fatigue, and/or inactivity. Low-cost commercial motion capture devices in combination with powerful computing devices that are capable of running sophisticated algorithms are becoming widely available for home use, technology being driven by the gaming industry. Our study has shown that estimating muscle forces that drive motion can enable a new type of automated assessment of age-related changes in movements and post-stroke motor deficits. This approach can be potentially useful as part of telemedicine and mobile health initiatives, where the patient’s health needs to be monitored remotely.

Most aged participants reached slower to maintain the accuracy specified by the virtual targets, demonstrating the classical speed-accuracy tradeoff (Fitts 1954; Young et al. 2009). This was reflected in mainly lower performance index in at least one and sometimes both arms in all but one aged participant. In the one aged participant (A5 59-year-old male) the performance index was higher than the mean of young controls in corresponding movements for both arms, and the movements were highly symmetrical even when quantified with MTd R^2^, which suggests that this participant did not have motor deficits but rather performed the movements using a different strategy. Two other aged participants (A3 and A4, both 58-year-old females) performed reaching movements less symmetrically with lower performance indices for their dominant right arms. The rest of the aged participants (male A1 and female A2, both 58-year-old) had symmetrical deficits so that their performance indices were lower in both arms. In A1 the bilateral reduction in performance index was due to the reduced speed of movement, while in A2 this was due to the increased variance of movement. The clustering analysis grouped participants A1 and A5 primarily with young controls, indicating that their sometimes slower (A1) and quirky (A5) movements were part of typical differences in how movements are produced by individuals rather than an indication of age-related motor deficits. The differences between aged and young participants were larger when measured from MTd compared to joint angle, indicating that the force-based measures are more sensitive to age-related motor deficits than motion-based measures are. Altogether, this shows that force-based metrics can be used to characterize with high sensitivity the individual differences in reaching movements associated with aging and distinguish them from typical inter-subject variability.

Here we show that force-based measurements of post-stroke motor deficits are more sensitive to individual pathology compared to motion-based measurements. In four participants with stroke (S3 and S7 with a stroke in medulla and S5 and S8 with a stroke in left and right MCA respectively, Table 1) the hemiparesis affected movements of both limbs as shown by reduced performance index for both limbs and reduced symmetricity of movements between limbs. In S4 with a lacunar infarct in posterior right putamen, the performance index of the ipsilesional arm was higher than that for aged participants and similar to that for young controls, indicating a different strategy for reaching movements, possibly similar to A5, rather than the presence of motor deficits in the ipsilesional arm. In S6 with a right MCA stroke the performance index for the ipsilesional arm was not different from that for aged nor for young participants, also indicating the absence of detectable motor deficits in the ipsilesional arm. However, the MTd R^2^ in both these participants was lower than in young controls, indicating that their movements were asymmetrical. In the last two participants with stroke (S1 with a left MCA stroke and S2 with a left caudate lenticular nucleus stroke), the effect of stroke on their reaching with either arm was very mild. Their reaching was slower, which reduced the performance indices for their contralesional arms. However, their movements were symmetrical with MTd R^2^ indistinguishable from young participants. The clustering analysis grouped participants S1, S2, and S6 primarily with young controls, indicating that their sometimes slower (S1 and S2) and asymmetrical (S6) movements were part of typical differences in how movements are produced by older individuals rather than an indication of motor deficits caused by stroke. The differences between participants with stroke and young controls were larger when measured from MTd compared to joint angle, indicating that the force-based measures of individual motor deficits due to stroke are much more sensitive than motion-based measures in quantifying the subtle multi-joint differences in complex 3D reaching movements. This also shows that the individual motor deficits caused by stroke can be distinguished from the typical differences in how movements are produced by individuals and from age-related changes in movement production.

The dynamic component of active muscle torque is indicative of intersegmental coordination, something that cannot be easily inferred from movement observation. Our results have shown that most Stroke group participants show a deficit in all metrics derived from MTd. This is consistent with published observations showing deficits in intersegmental coordination and the phasic muscle activity patterns after a stroke during the performance of planar reaching tasks and some unconstrained reaching tasks (Bastian et al. 1996; Beer et al. 2000; Cirstea et al. 2003; Levin 1996; Manto and Bosse 2003). Furthermore, the deficits measured with MTd were generally present for all movement directions (Fig. 3, 4C, and 5B), indicating that the disruption in intersegmental coordination caused by stroke is general and can be observed in most reaching movements regardless of the starting position, speed, and direction of reaching. This shows for the first time that the deficits in intersegmental coordination caused by stroke can be quantified in unconstrained 3D reaching movements while controlling for posture-dependent forces and other sources of inter-subject variability. This method for quantifying intersegmental coordination may help increase the efficacy of robotic or robot-assisted post-stroke rehabilitation (Patton et al. 2006; Sanchez et al. 2006; Volpe et al. 1999, 2000, 2001). New approaches for designing the pattern of robot assistance or resistance to appropriately titrate the difficulty of therapy while avoiding moving the arm passively, are needed (Kahn et al. 2014; Lum et al. 2002). Quantifying deficits in the dynamic component of active muscle torque and setting robotic devices to assist or resist only this component may be a way to personalize and standardize intervention and improve the recovery of intersegmental coordination.

Furthermore, quantifying the dynamic component of muscle torque while controlling for the gravity-compensating component enables direct comparison between unassisted and unconstrained 3D movement and movements performed with support against gravity or other external assistance or resistance. In particular, supported planar 2D reaching tasks are now used more widely for robot assisted therapy in combination with quantitative metrics derived from the same planar tasks (Coderre et al. 2010; Kwakkel et al. 2019; Scott and Norman 2003). However, these movements are constrained and not overtly functional, which makes it difficult to quantify the carry over effect from therapy based on planar movements into real-world effects on 3D functional movements in a meaningful way that helps inform future interventions. The method used here to derive quantitative assessment metrics from the individual dynamic component of muscle torque can help bridge the gap between evidence from studies of constrained or robotically manipulated movements and research with functional and unconstrained movements.

## Acknowledgments

We thank the study participants for generously giving their time, and M. Powers for her assistance with participant recruitment and the general support of the study; we acknowledge the technical support expertly provided by B. Pollard; we are thankful for insightful and critical comments of S. Yakovenko.

